# Study of the AMP-activated protein kinase role in energy metabolism changes during the postmortem aging of yak *longissimus lumborum*

**DOI:** 10.1101/670851

**Authors:** Yayuan Yang, Ling Han, Qunli Yu, Yongfang Gao, Rende Song

## Abstract

To explore the postmortem physiological mechanism of muscle, activity of adenosine monophosphate activated protein kinase (AMPK) as well as its role in energy metabolism of postmortem yaks were studied. In this experiment, we injected 5-amino-1-beta-d-furanonyl imidazole-4-formamide (AICAR), a specific activator of AMPK, and the specific AMPK inhibitor STO-609, to observe the changes in glycolysis, energy metabolism, AMPK activity and AMPK gene expression (PRKA1 and PRKA2) in postmortem yaks during maturation. The results showed that AICAR could increase the expression of the PRKKA1 and PRKAA2 genes, activate AMPK and increase its activity. The effects of AICAR include a lower concentration of ATP, an increase in AMP production, an acceleration of glycolysis, an increase in the lactic acid concentration, and a decrease in the pH value. In contrast, STO-609 had the opposite effect. Under hypoxic adaptation, the activity of the meat AMPK increased, which accelerated glycolysis and metabolism, and more effectively regulated energy production.

## 1. Introduction

Yaks adapted to high altitudes because of the colder climate. This process included the maintenance of the production of adenosine triphosphate (ATP) through an increase in glycolysis. Studies have shown that yaks have specific metabolic mechanisms that enable them to adapt to a hypoxic environment to attain an adequate supply of energy and a demand balance under hypoxic conditions. (Zuo et al., 2017; Hardie, Ross, Hawley, 2012; Hardie, 2003). AMP-activated protein kinase (AMPK), as an important cellular energy sensor, is critical for the regulation of the metabolism of energy and the subsequent quality of the meat. Under hypoxic conditions, the body is under stress; metabolism is strengthened; ATP consumption is increased; the ATP concentration is decreased; the AMP production is increased, and a high concentration of 5’-AMP and AMPK gamma subunits interact to activate AMPK. Ding et al. (2014) studied the activity of lactate dehydrogenase (LDH) in yaks at three different altitudes, and its activity positively correlated with altitude. LDH is the key enzyme for anaerobic glycolysis, indicating that yaks at higher altitudes are more dependent on energy metabolism (Chen, 2003; Wojtaszewski, 2000; Musi, 2001; Park, 2002)

Under hypoxic conditions, the body is under stress; metabolism is strengthened; ATP consumption is increased; the ATP concentration is decreased; AMP production is increased, and a high concentration of 5’-AMP and AMPK gamma subunits interact to activate AMPK (Park et al., 2002). Research on the activity of lactate dehydrogenase (LDH) in yaks at three different altitudes indicated that the enzyme is positively correlated with altitude. LDH is the key enzyme for anaerobic glycolysis, indicating that yaks at higher altitudes are more dependent on the metabolism of energy (Ding et al., 2014). Thus, it is necessarily to additionally study changes of energy metabolism and AMPK activity in a hypoxic environment.

The enzyme AMPK is a heterotrimer consisting of α, β, and γ subunits. Its primary role is thought to be the critical regulation of energy metabolism (Hardie, 2004; Carling, 2004; Winder, 2001; Hardie & Carling, 1997). AMP/ATP ratio increase in muscle cells is thought to result in the activation of AMPK. This activation results in the phosphorylation of AMPK at Thr172 by a kinase that remains unidentified. Following its activation, AMPK activates glycogenolysis/glycolysis and consuming/catabolic pathways that generate ATP (Scott, Pan, & Hudson, 2003; Minokoshi et al., 2004; Hardie, Kemp et al., 1999; Hardie & Carling, 1997; Corton, Gillespie, & Hardie, 1994). Any type of cellular stress can cause AMPK activation. Physiological AMP/ADP elevation occurs as a result of stress, such as low nutrients or prolonged exercise. As previous studies have found, the initiation of glycolysis in ischemic heart AMPK activation plays an important role (Sambandam & Lopaschuk, 2003). Thus, the data accumulated confirms that hypoxia is a characteristic of post-mortem skeletal muscle and ischemic heart disease. Therefore, post-mortem glycolysis may be regulated by AMPK.

Previous research demonstrated the activation of AMPK in pork loins, which develop into PSE meat. This finding suggests that a key role of AMPK in regulation of postmortem glycolysis (Shen, Means, Underwood, et al., 2006). Therefore, the role of AMPK may be the regulation of glycolysis in post-mortem skeletal muscle. If so, the enzyme may be a logical target to manipulate to intervene in the process of PSE development and cause its reduction, since AMPK activity depends on the postmortem skeletal muscle pH values. Therefore, we further studied AMPK role in muscle glycolysis regulation in post-mortem meat, using specific AMPK activators and inhibitors to detect whether the induction of AMPK by 5-amino-1-β-D-ribofuranosyl-imidazole-4-carboxamide (AICAR) and STO-609 affect post-mortem muscle glycolysis. Recent research in rat skeletal muscle used a cell-permeable compound AICAR to activate AMPK to study its possible task in controlling glucose metabolism in this tissue (Fan et al., 2009; Pold et al., 2005; Thomson et al., 2009). These studies involved administration of either *in vivo* or in *vitro* AICAR to skeletal muscle for varying amounts of time. In addition, changes in carbohydrate metabolism were accessed using different methods. These methods included a muscle preparation that had been isolated and incubated, a hindquarter preparation that had been perfused, or tissues analyses following a euglycemic clamp or treatment. To our knowledge, the effect of reactive AICAR on the AMP-activated protein kinase of mice longissimus lumborum has only been examined in one study. Among other studies, we found that injecting a dose of 250 mg/kg AICAR had no effect on the glycogen binding in the diaphragm (respiratory muscle) of mice fed or fasting (Vincent et al., 1996). At the same time, the effects on glucose transport in yak skeletal muscle due to AICAR treatment remain unclear. In comparison, the inhibitory effect of STO-609, an AMPK inhibitor, was used to inhibit food intake and therefore weight gain in mammals (Hayes et al., 2009; Kim et al., 2004). This further clarifies the effects of AMPK in the demodulation of postmortem glycolysis.

This study was performed to analyze AICAR and STO-609 influences on AMPK gene (PRKAA1, PRKAA2) mRNA expression and AMPK activity on glycolysis and energy metabolism during post-mortem of yak after slaughter. It laid a foundation for the establishment of the theoretical system of energy metabolism in postmortem yak meat.

## 2. Methods and Materials

### 2.1. Animal treatment

The *M. longissimus lumborum* is the LL, the 12^th^ rib that is anterior to the last lumbar vertebrae, and they were randomly extracted from a slaughterhouse (Yushu Tibetan Autonomous Prefecture, Qinghai Province, China). Ten Qinghai yak bulls that were the same age (36-38 months old) and weighed 241-280 kg were fed the same diet from the same batch. Each yak was tested. The ribs were immediately frozen in liquid nitrogen as the sample for 0 h. The remaining amount of each 40 g aliquot of the muscle pieces was divided into two portions. One portion was treated as the control, while the other was treated with injections of a 1:1 ratio (w/v) of 10 mM AICAR (Sigma A9978) and STO-609 (Sigma Aldrich). All samples were subsequently stored at 4 °C for 12, 24, 72, 120, and 168 h. All samples collected at these times were stored at −80 °C until further use.

### 2.2 Measurement of the pH

The pH values of all the loins at each time point were measured using a Testo® 230 meter (Lenzkirch, Germany). pH values were measured using pH meter calibrated standard buffer solutions with pH values equal to 4.0 and 7.0 (Mallinckrodt Chemicals, USA). Prior to calibration, buffers solutions were stored at 20 °C according to procedure mentioned elsewhere (Fernandez, Neyraud, Astruc, & Sante, 2002).

### 2.3 Lactic acid concentration

A total of 500 mg of frozen muscle samples was homogenized using 500 mL of 0.9% saline and then centrifuged at 4200*g* at 4 °C for 10 min. Following a 50-fold dilution of the supernatant, standard commercial kits from Blue Gene Biotech Co. (China) were used to measure the lactic acid contents. The optical density (OD_450_) was measured immediately using an ELISA microplate reader. The values measured at each concentration of the standard were used to prepare a calibration curve (Shen, Means, Thompson, et al., 2006).

### 2.4 ATP, ADP, AMP and IMP activity

In accordance with Hou’s method (2011), approximately 3 g of frozen muscle was centrifuged for 10 min at 15,000 × *g* (Heraeus, Biofuge fresco, Germany) at 4°C. The supernatant was mixed with 1.44 mL of 0.85 M K_2_CO_3_ and filtered through a 0.2 μm membrane. The ATP, ADP, AMP and IMP contents were analyzed using Agilent 1100 Chromatography at 254 nm detection wavelength. A reversed phase C18 column was used, and the flow rate was 1 mL / min. Quantitative analysis was conducted on the basis of retention time and peak area.

### 2.5 AMPK activity

AMPK activity measurements were based on AMPK specific phosphorylation of a SAMS peptide (Shen et al., 2006). Briefly, SAMS peptide substrate (His-Met-Arg-Ser-Ala-Met-Ser-Gly-Leu-His-Leu-Val-Lys-Arg-Arg, obtained from Invitrogen, USA) was used for the assay. As-obtained muscle homogenate was centrifuged at 13000*g* at 4 °C for 5 min. 10 μL of supernatant was incubated for 10 min at 37 °C at pH 7.0. Its final volume was 50 μL, and it contained 0.2 mM of ATP ^+ 2^ μCi [^32^P] ATP, 0.2 of mM AMP, 5 mM of MgCl_2_, 0.2 of mM SAMS peptide, 80 of mM NaCl, 0.8 mM of dithiothreitol, 0.8 mM of EDTA, 8% (w/v)of glycerol, and 40 mM of 4-2-hydroxyethyl-1-piperazineethanesulfonic acid. 20 μl of this mixtures was removed and placed on Whatman P81 filter paper (Whatman, England) that had been cut into 2 cm × 2 cm pieces. Six washes of 1% phosphoric acid were conducted to remove the ATP. Finally, the filter paper was immersed in 3 mL of Scinti Verse (obtained from Fisher Scientific, USA). AMPK nanomolar peptide activity phosphorylation was expressed per minute per gram of muscle.

### 2.6 Immunoblotting

AMPK was analyzed using the frozen yak LL muscle derived from these methods as previously described (Shen et al. 2005). Briefly, 0.05 g of muscle was homogenized at top speed for 10 s on ice using a Polytron homogenizer (IKA Works, Inc., Wilmington, NC). Five hundred milliliters of precooled buffer was used to homogenize the tissue. The buffer contained 20 mM of Tris–HCl with pH value equal to 7.4 and at initial temperature equal to 4 °C as well as 2% SDS, 5 mM EGTA, 5 mM EDTA, 1 mM DTT, 100 mM NaF, 2 mM sodium vanadate, 10 mg/ml pepstatin, 0.5 mM phenylmethylsulfonyl fluoride (PMSF) and 10 mg/ml leupeptin (Veiseth, Shackelford, Wheeler, & Koohmaraie, 2001; Raser, Posner, & Wang, 1995). Each of the muscle homogenate was mixed with an equal volume of 2x SDS-PAGE loading buffer containing 0.5 M TrisHCl (pH 6.8), 2% (v/v) 2-mercaptoethanol, 20 vol% glycerol, 4.4% (w/v) SDS, and 0.01% bromophenol blue (boiled for 5 min prior to electrophoresis).

The gels were cast using a BioRad mini-gel system (Richmond, CA, USA) that was also used to perform the SDS-PAGE electrophoresis. Gradient gels of 5%-20% were used to separate the proteins. When the electrophoresis was complete, proteins separated on the gels were moved to nitrocellulose membranes using buffer that contained 20 mM Tris-base, 0.1% SDS, 20% methanol, and 192 mM glycine. Next, the membranes were subjected to incubation in a blocking solution comprised of 5% zero fat dry milk in TBS/T (150 mM NaCl, 50 mM Tris–HCl (pH 7.6), and 0.1% Tween-20 for 1 h. Them these membranes were subsequently incubated overnight for the western blotting using one of two types of antibodies: monoclonal anti-β-actin antibody (Sigma-Aldrich, USA) or a primary antibody, anti-phospho-AMPKα (Thr 172, obtained from Cell Signaling Technology, USA). The membranes were washed 3 times (5 min each) using 20 ml of TBS/T following incubation with the primary antibody. The next step involved the incubation of the membranes with horseradish peroxidase-conjugated secondary antibodies that had been diluted 5-fold. These membranes were agitated gently for 1 h in TBS/T, followed by washing three times (10 min each time). ECL western blotting reagent (from Amersham Bioscience) was used to visualize the membranes by exposing them to MR film (Kodak, Rochester, NY). An Imager Scanner II and Image Quant TL software were used to quantify density of the bands (Shen et al., 2005). Samples obtained after all these treatments were analyzed on a single gel to decrease variation between the blots. Reference band density was used to normalize band densities from different blots. In addition, the density of the ß-actin band was also used to normalize the band densities.

### 2.7 Real-time PCR analysis

Real-time reverse transcription (RT)-PCR was used to quantify expression levels of the genes selected for the analysis (Monika et al., 2006). Briefly, TRIzol reagent (Invitrogen Corp., USA) was used to extract total RNA from the LL based on method recommended by a manufacturer. RT was performed using Oligo(dT) random 6-mer primers from a Prime Script RT Master Mix kit (TaKaRa, Dalian, China) according to the manufacturer’s instructions. A SYBR Premix Ex Taq kit (TaKaRa, Dalian, China) was used to perform quantitative PCR on a CFX96 Real-Time PCR detection system (BioRad). All of the experiments analyzed each RNA sample in triplicate. In addition, each experiment involved a negative control that lacked a cDNA template. ΔΔCt method (based on report of Livak & Schmittgen, 2001) was implemented to obtain relative expression levels of the target mRNAs.

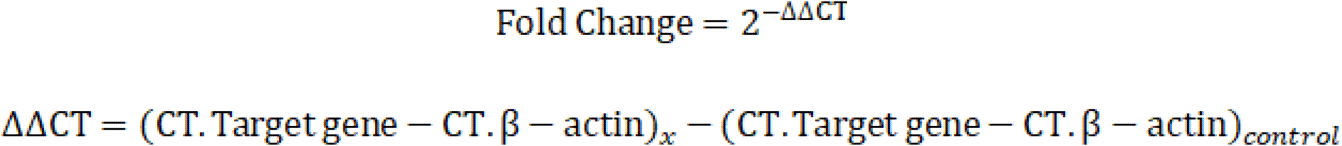

### 2.8 Data processing and statistical analyses

A one-way analysis of variance (ANOVA) was implemented to obtain statistical significance for the differences using IBM SPSS 19.0 Software (SPSS, Inc., Chicago, IL, USA). Duncan’s multiple range test was used for significance determination among the groups. At *P* < 0.05, results were considered statistically significant. The dynamics and graph plotting were conducted using Origin 8.0 software. Each experiment was repeated at least three times.

## 3. Results

### 3.1 pH value determination

AICAR injection in the LL increased the pH decline, while STO-609 injection in the LL decreased the pH decline in the postmortem yak LL muscle (Figs. 1). At 0 h postmortem, no difference in muscle pH was detected between these three treatments. However, at 12 h postmortem and through the remaining sampling times, the muscle pH of the control yak was higher (*P* < 0.05) than that of the AICAR-treated yak but lower than STO-609-treated yak. At 24 h postmortem, the pH of the AICAR-injected yak muscle remained less than 6.0, showing that the glycolytic rate was strongly activated (Fig. 1).

**Fig. 1.**
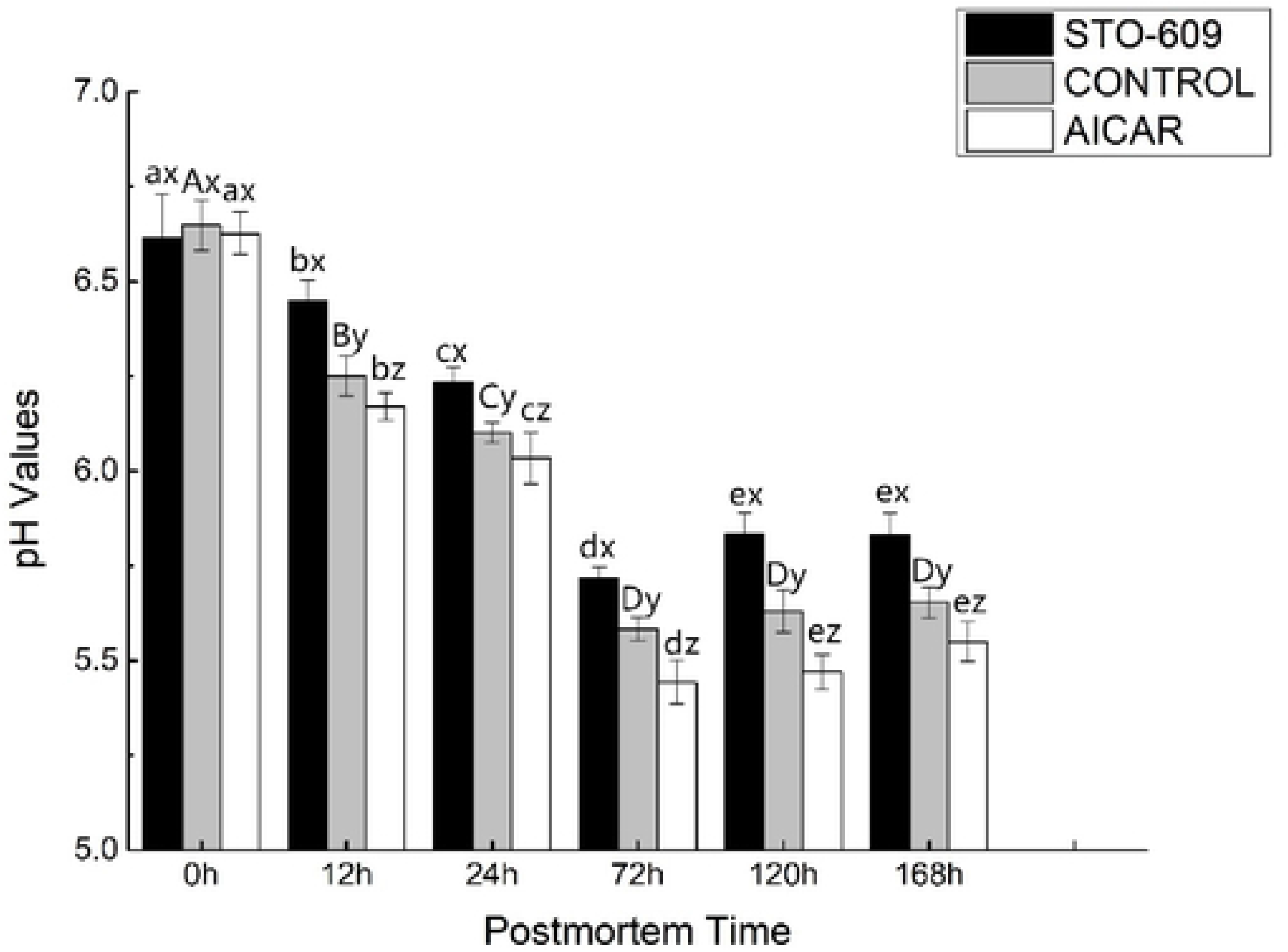
pH values of postmortem yak LL muscle with AICAR and STO-609 treatment. One-way ANOVA was used for statistical analyses between the control group and two treatment group at 0 h to 168 h (x, y, z *P* < 0.05). Duncan’s New Multiple-range test was used for the differences between the control group and two treatment group at 0 h to 168 h. At 0 h, the lowercase letters represent the difference of the treatment group and the capital letters represent stands the difference of the control group over time (*P* < 0.05). Error bars indicate the standard errors of the mean.

### 3.2 Lactic acid concentration

Increased glycolysis in skeletal muscle of yak injected with AICAR was confirmed by increased lactic acid accumulation rate (Fig. 2). In addition to lowering the pH, the injection STO-609 after reduction yak LL mortem muscle lactate accumulation (Fig. 2). STO-609 injection yak postmortem glycolysis sarcopenia, decreased lactate accumulation rate. After 0 h, muscle lactic acid concentration between the three treatments difference was not detected. From 0 to 72 h postmortem, the lactic acid concentration increased 126.56 ± 5.89 mg/g muscle in control group and only 96.32 ± 3.19 mg/g muscle in the STO-609 group, respectively, which indicates that STO-609 injection inhibited lactic acid production in postmortem muscle at the initial stage. Between 0 and 72 h, the lactic acid concentration in the AICAR group increased by 132.51±6.32 mg/g. This result indicates that AICAR injection activates lactic acid production in the initial stage of post-mortem muscle. Results confirmed that AICAR promotes muscle glucose uptake in yak but STO-609 inhibited lactic acid production and suggested that this process may be mediated by a novel glucose transporter.

**Fig. 2.**
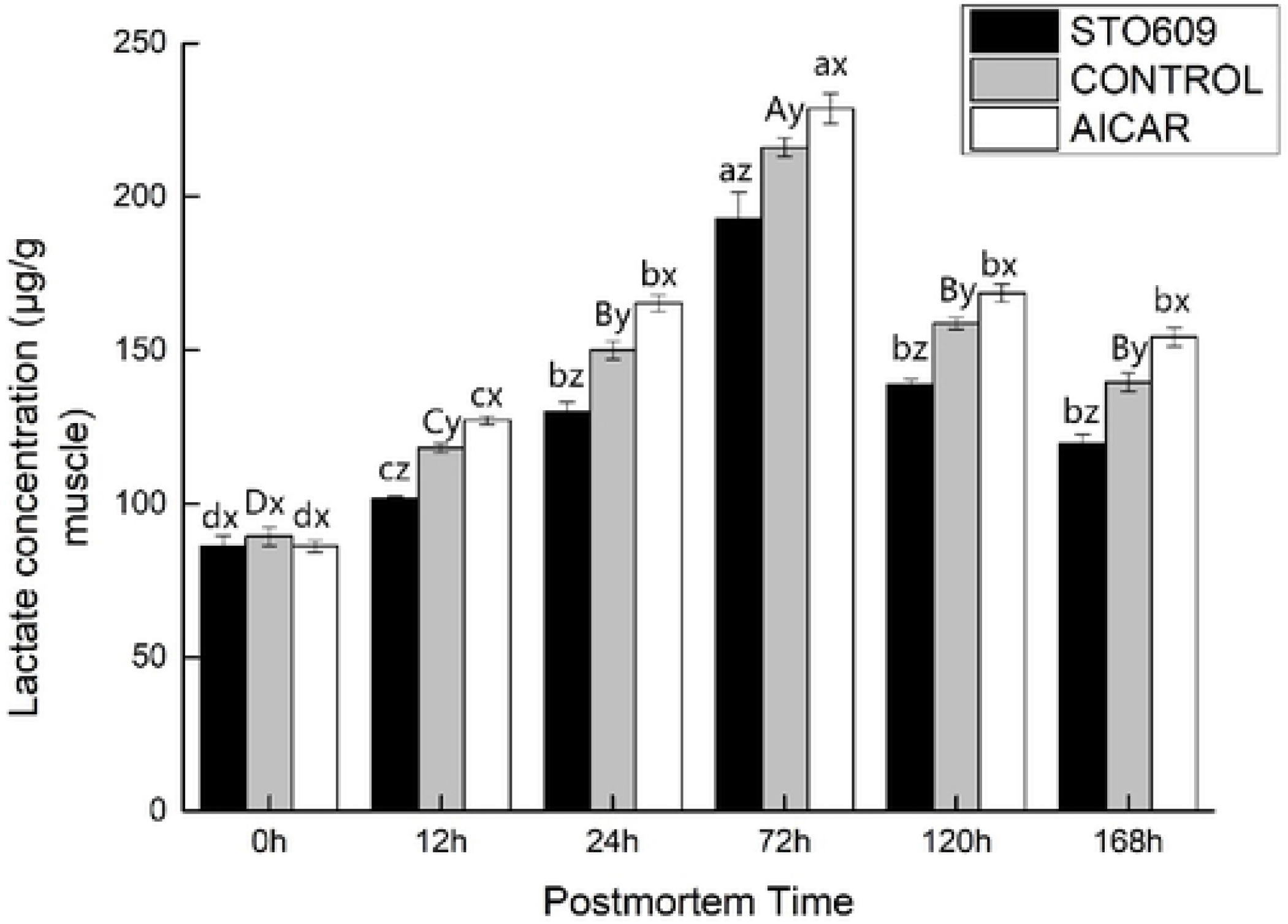
Lactic acid in postmortem yak LL muscle with AICAR and STO-609 treatment.

### 3.3 ATP, ADP, AMP and IMP activities

The nucleotide concentration in the yak LL muscle was measured in this study. After the 0 h control, there were no significant differences the in nucleotide concentration observed between the control and treatment groups (Table 2). However, AICAR injection increased the ATP levels of the skeletal muscle, while decreasing the concentrations of AMP and IMP (*P* < 0.05). This result could be due to the inhibition of glycolysis in the LL muscle by the AICAR injection (Table 2). The AICAR injection inhibited glycolysis in the postmortem muscle (Figures 2, 3). As a result of this inhibition, less ATP was produced, and the ATP concentration increased within 12 hours following death (Table 2). The IMP in the muscles of the yaks was also significantly higher than the AMP following the slaughter of the animals, and these results are consistent with those of previous studies (Shen et al., 2007; Shen et al., 2006). This result reiterated an observation that postmortem skeletal muscle delamination results in the rapid conversion of AMP to IMP. At the 0 h control, the nucleotide concentrations between the three groups did not differ significantly (Table 2). However, STO-609 injection resulted in a decrease in the concentration of ATP in the skeletal muscle and an increase in AMP and IMP at 12 h postmortem (*P* < 0.05). This result can be explained by the inhibition of glycolysis due to the injection of STO-609 into the yak LL muscle. STO-609 injection inhibited glycolysis of the postmortem muscles (Figures 1, 2), resulting in lower amounts of ATP production and therefore, a decrease in ATP concentration 12 h (Table 2). In addition, the data showed that the levels of IMP greatly exceeded those of AMP in the postmortem yak muscles, buttressing the results of previous reports (e.g. Shen et al., 2007; Shen, Thompson et al., 2006). This finding reiterates the importance of AMP. The skeletal postmortem muscle undergoes rapid conversion to IMP by the process of deamination. A decrease in the glycolysis of yak LL muscle injected with STO-609 indicates that the glycolysis of the skeletal postmortem muscle is partially regulated by AMPK.

**Table 1.**
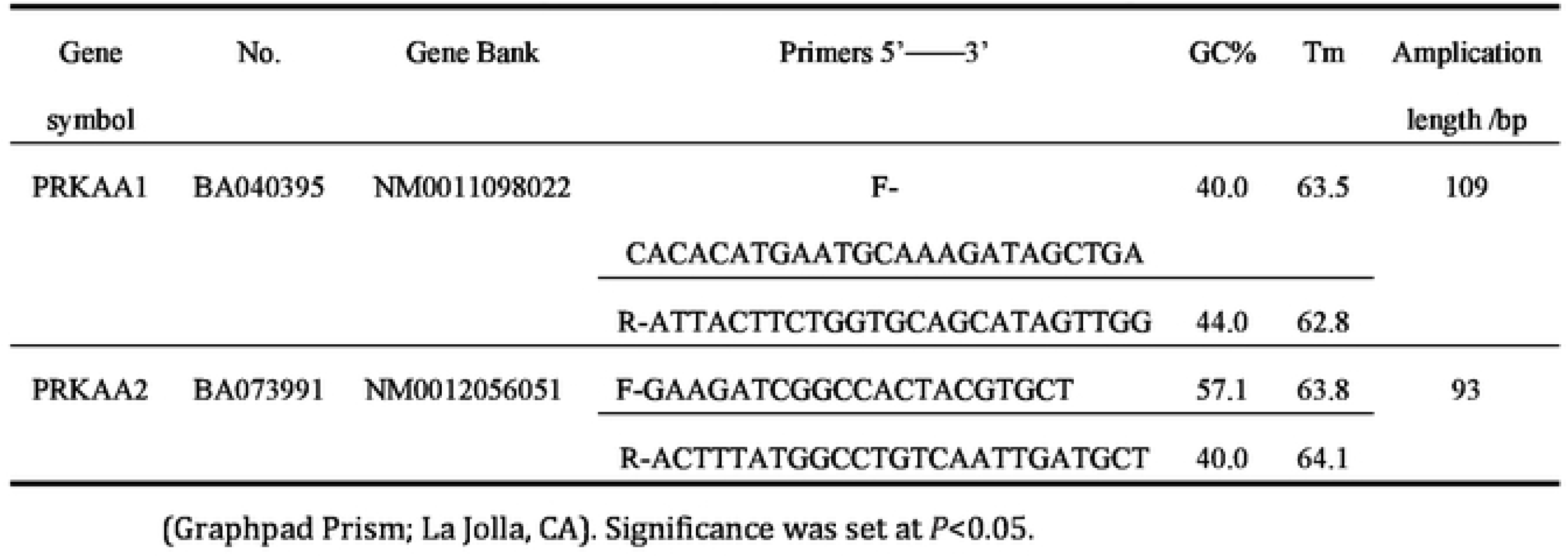
Primer sequences and parameter used for real-time quantitative PCR.

**Table 2.**
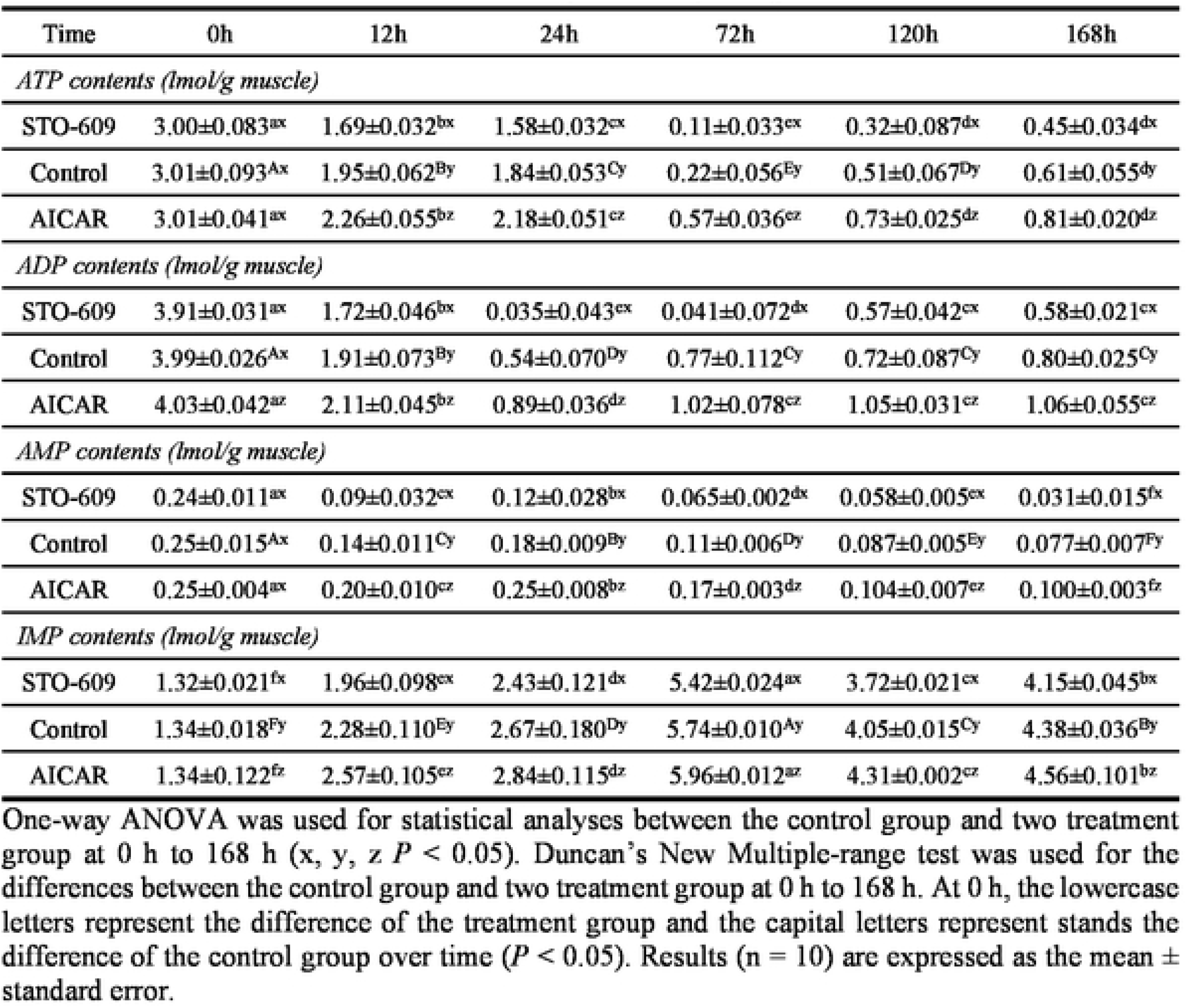
Nucleotide concentration in postmortem yak *longissimus dorsi* muscle.

**Fig. 3.**
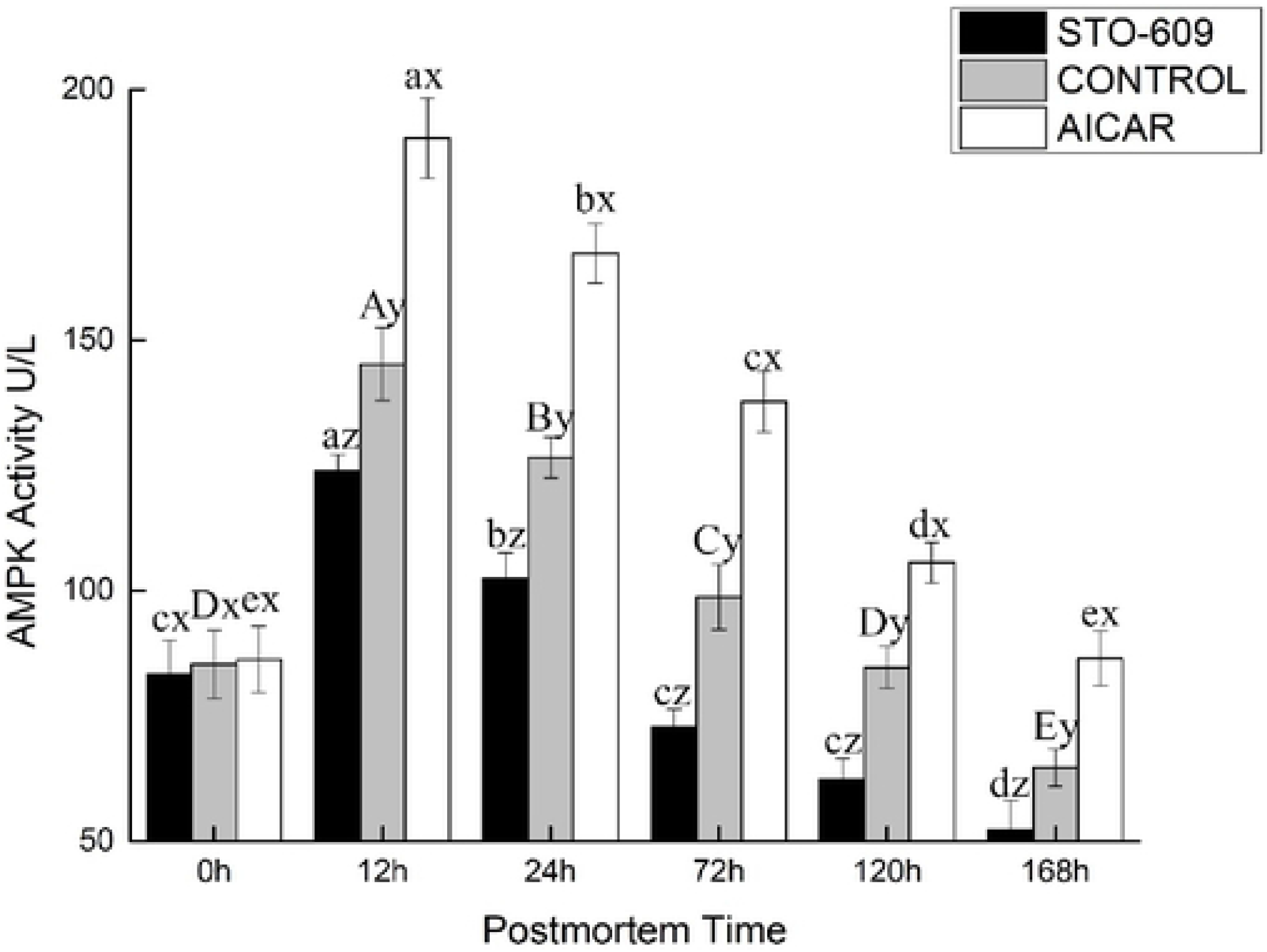
AMPK Activity of postmortem yak LL muscle with AICAR and STO-609 treatment. At a specific postmortem time, indicates significant difference at *P* < 0.05.

### 3.4 AMPK activity

Figure 3 shows the activity of AMPK in the postmortem yak LL. At the 0 h postmortem control, the activities of AMPK were 1.56 ± 0.06, 1.19 ± 0.13, and 1.00 ± 0.07 nmol of ATP per min per gram of the muscle mass for the AICAR and STO-609 treatments and the control, respectively. Representative AMPK activity is shown in Figure 3. The activation of AMPK was more rapid in the AICAR group and reached its maximal level at 12 h postmortem (Figure 3). At this same time, AMPK activity in the AICAR treatment exceeded those of the control and the STO-609 groups at 2.39 ± 0.19 nmol of ATP per min per gram of the muscle mass. These results indicated that a more rapid activation of AMPK, and therefore higher activity, explained the faster decline in pH values and the higher rate of glycolysis during the early stage postmortem muscle of the yak.

### 3.5 Immunoprecipitation of AMPK

The main goal of this work was to study influence of STO-609 and AICAR on postmortem glycolysis and AMPK activity. STO-609 functions as a specific competitive inhibitor of AMPK and has been well studied (Kim Et al., 2004; Blattler, Rencurel, Kaufmann, & Meyer, 2007). Representative immunoblot shown in Figure 5 demonstrates AMPK phosphorylation. As expected, the postmortem muscle yak LL AMPK phosphorylation was reduced following the injection of STO-609 (Figure 5). The activities of AMPK were higher in the skeletal muscle after death following STO-609 without sputum injection, and the greatest amount of activity was detected at 12 h postmortem. However, in the samples treated with STO-609, no significant change in the levels of AMPK activity as a result of the aging time was observed (Figure 4). After 0-12 h postmortem, the level of phosphorylation of AMPK in the control group increased by 0.39 ± 0.02 arbitrary units, and this was greater than the 0.20 ± 0.01 arbitrary units in the LL muscle after the STO-609 injection (*P* < 0.05). In the muscles of the yak injected with STO-609, AMPK phosphorylation was decreased because the subunit Thr 172 delayed the decrease in AMPK activity. However, from the 0 to 12 h samples postmortem, the phosphorylation of AMPK increased by 0.39 ± 0.02 arbitrary AICAR units. This increase was higher than the 0.20 ± 0.01 arbitrary units in the postmortem LL yak muscle without injection (*P* < 0.05). Increased AMPK phosphorylation of the subunit at Thr 172 indicates higher AMPK activity in yak postmortem muscle subjected to AICAR injection.

**Fig. 4.**
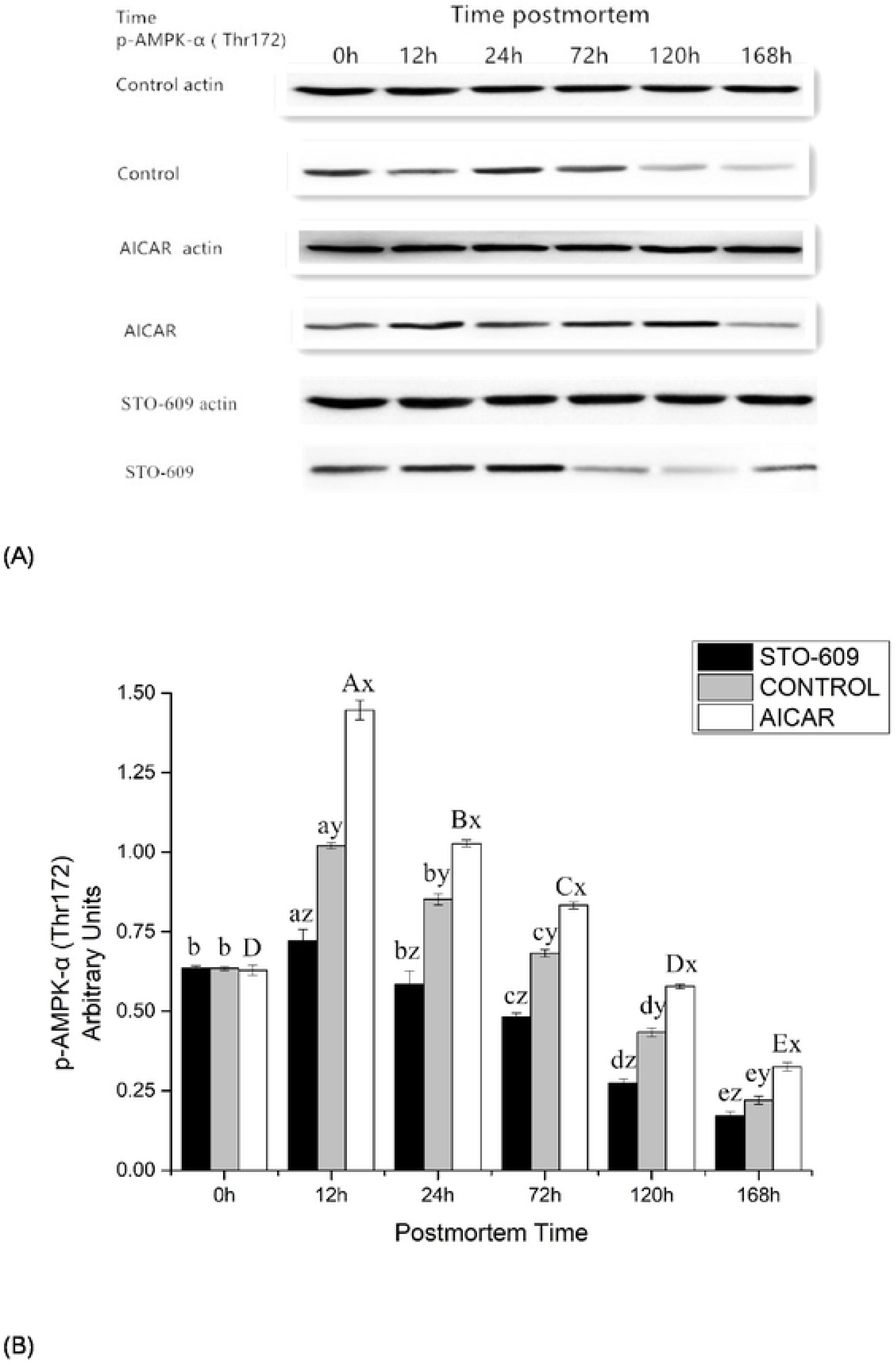
(A) Effects of yak LL muscle injection of AICAR and STO-609 on AMPK phosphorylation (Thr 172) in postmortem yak LL muscle. Representative immunoblots of AMPK phosphorylation and b-actin, and the relative band density of phospho-AMPK after normalizing to b-actin were shown. (B) Densitometric analysis of AMPK expression of bovine muscle during postmortem aging.

**Fig. 5.**
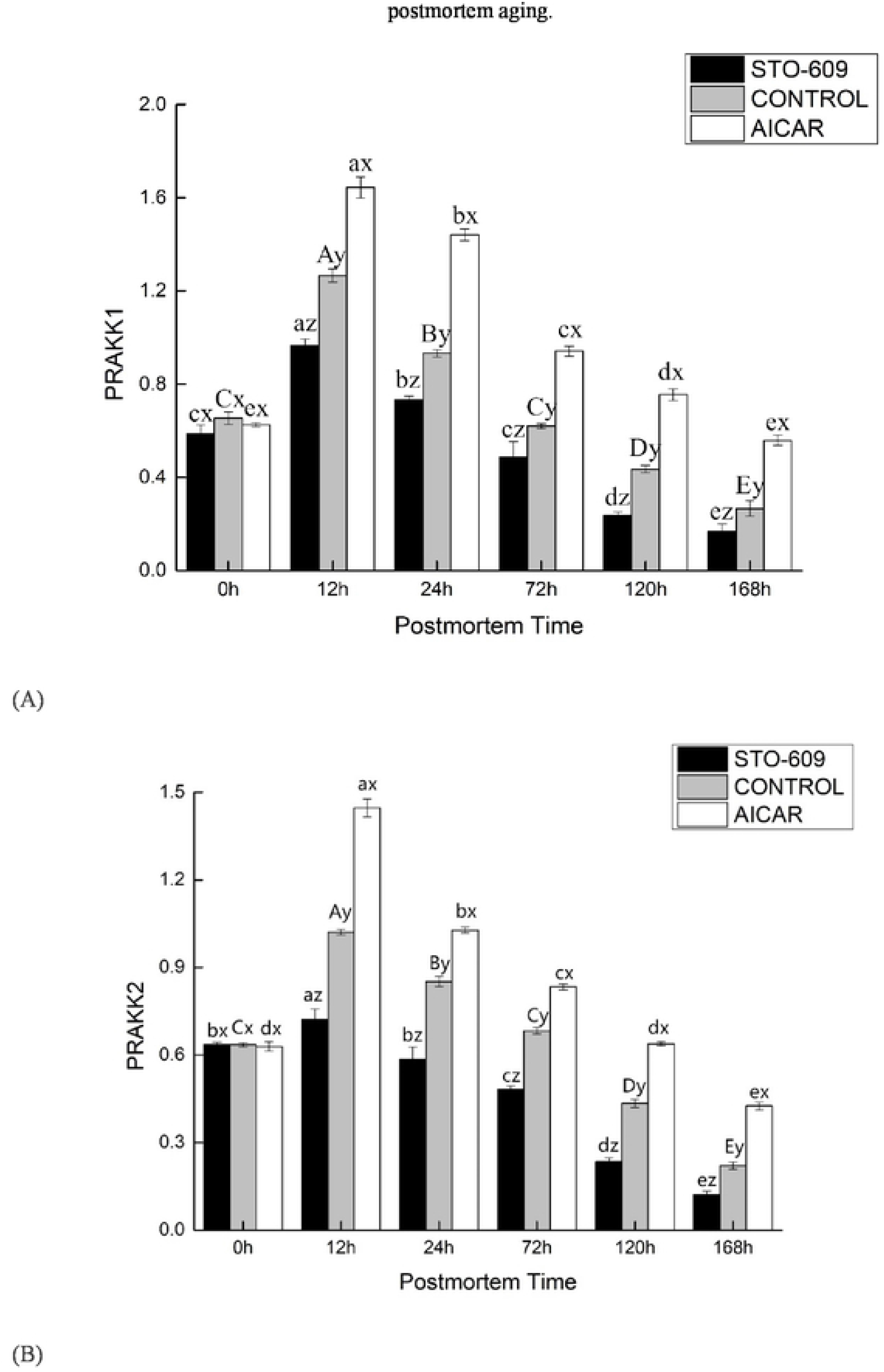
Effect of AICAR and STO-609 on AMPKα1(A), AMPKα2 (B) mRNA were treated with or without AICAR (10 mM) and STO-609 (10 mM) for the time as indicated above and total RNA was subjected to real-time RT-PCR as described in Materials and methods. The results were expressed as a relative value compared to the untreated sample as 100%. All data were represented as means ± SEM of three independent experiments. *P* < 0.05, compared with the untreated control.

### 3.6 Gene Expression of AMPK

Previous studies demonstrated that line injections of AICAR and STO-609 in yak could involve the hypothalamic AMPK system. To determine whether the AMPK system mediated the effect of AICAR, the gene expression of different AMPK subunit levels of gene expression was tested. The results indicated no mRNA level increase of the yak hypothalamic α1 subunit of AMPK after AICAR injection The phosphorylation of AMPK is mediated by the cooperation of these different subunits, which was shown to correlate with AMPK activity. This study indicates that the AICAR injection caused different effects on the catalytic and regulatory AMPK subunits. The mRNA expression of the regulatory subunits α1 and α2 was stimulated by AMPK. However, this was not the case for the phosphorylation or mRNA expression of catalytic subunit α. These results suggest the increase in AMPK activity may be due to an independent mechanism that could be inhibited by STO-609. The fact that the regulatory subunits β and γ can only result in catalytic activity when they are in a complex with the α subunits has strong implications for the role of the catalytic subunits in the activity of AMPK. It suggests that their availability is a key determinant of this activity. Although there is not enough data on AMPK role in glucose metabolism regulation to make firm conclusions, our research provides the first analysis of the effect that an AMPK agonist has on glucose metabolism in yaks. These results confirm the promotion of glucose uptake in yak muscle by AICAR. In addition, the results suggest that this process may be due to the mediation of a novel glucose transporter. In total, these results provide confirmation that STO-609 acts as a potent AMPK inhibitor and causes a reduction of AMPK activity in postmortem yak muscles.

## 4. Discussion

This study focused on how AICAR and STO-609 affected the AMPK pathway, which is related to energy metabolism in the yak LL muscle. Our results suggest that AICAR and STO-609 induce changes in parameters involved in energy metabolism. AMPK functions to regulate the metabolism of energy and substrates, primarily the metabolism of carbohydrates and the homeostasis of whole-body energy. AMPK acts as an energy sensor for the whole body to meet both body energy and cellular requirements by integrating different signaling pathways, while also activating energy-producing processes and inhibiting those processes that consume energy (Copenhafer, Richert, Schinckel, Grant, & Gerrard, 2006). AMPK primarily promotes fatty acid and glucose catabolism, while preventing the synthesis of glycogen.

It has been clearly proven that PSE development meat is because of its influence on rapid postmortem glycolysis that results in the early buildup of lactic acid (Solomon et al., 1998; Schwagele et al., 1994; Briskey & Wismer-Pedersen, 1961). Muscle pH values, which became PSE meat, declines at different rates than the values of normal muscle tissue; the pH values decline much more quickly (with 1.04 u/h speed) in the muscle that will be PSE meat in comparison to normal muscle (which have a speed equal to 0.65 u/h) at 37 °C (Bendall et al., 1963). The involvement of AMPK in both lipid and glucose metabolism has been reported. Stress induced by nutrients or exercise increases the level of AMP at the cellular levels. This increase accelerates beta oxidation of fatty acids as well as glucose transport into skeletal muscles In contrast, AMPK activation inhibits the several gene transcription, apoptosis as well as cholesterol and fatty acid syntheses. The injection of STO-609 reduces the decline in pH and the accumulation of lactic acid in postmortem murine LL muscle (Figures 1 and 2). In concert, the intraperitoneal injection of STO-609 into murine early stage postmortem muscle tissue demonstrated the initial inhibition of the level of postmortem glycolysis due to the development of lower lactic acid concentrations and higher pH values in the tissue. The injection of AICAR increased the accumulation of lactic acid and the decline in pH values in the muscle of postmortem murine LL (Figures 1, 2). In concert, the development of higher concentrations of lactic acid and lower pH values in the early stage yak postmortem muscle following AICAR injection indicated that the level of initial postmortem glycolysis increased due to the effects of AICAR. Implementation of combination of STO-609 as a specific AMPK inhibitor and commonly used AICAR as an activator of AMPK in an array of cellular systems suggest that AMPK mediates postmortem glycolysis reduction. Thus, this study confirms previous ones that indicate postmortem skeletal muscle glycolysis is AMPK-regulated (Shen et al., 2006; Shen & Du, 2005b).

AMPK performs an important part in substrate and energy metabolism due to its regulation of signaling pathways. AMPK monitors the availability of nutrients as well as AMP/ATP and ADP/ATP ratios, which enables it to sense the status of cellular energy levels. AMPK also regulates events in the cell by both inhibiting reactions and activates consuming ATP (e.g. protein and fatty acid syntheses) as well as cellular processes generating ATP (e.g. fatty acid oxidation, glucose uptake and glycolysis) (Hardie et al. 2006).

Previous reports demonstrated that preslaughter stress quickened depletion of muscle ATP levels. This reduction in the status of energy therefore results in the early and rapid activation of AMPK in early postmortem stages, which makes it more likely that pork loin will develop into PSE meat (Shen et al. 2007; Shen et al., 2006). Primary AMPK function is regulation of the internal cellular energy balance (Hardie et al., 2003; Hardie et al., 1999; Winder & Hardie, 1999). ATP depletion results in the activation of AMPK, or to be more precise, an AMP/ATP ration increase (Corton et al., 1994; Birnbaum, 2005; Witters et al., 2006). AMPK is affected by the binding of AMP occurring at low intracellular energy levels and high concentrations of AMP. AMPK changes its conformation and becomes a more effective substrate for LKB1, which is also known as upstream AMPK kinase (Hawley et al., 1995). Additionally, LKB1 activates AMPK by phosphorylating it (Woods et al., 2003; Shaw et al., 2004; WoodsHawley et al., 2003). Activation of AMPK results in the switching on of fatty acid oxidation and glycolysis, which results in the production of greater amounts of ATP inside the cells (Carling et al., 1987; Marsin et al., 2002; Carlson & Kim, 1973). Earlier research demonstrated that the halothane gene and the stress of the pre-slaughter process accelerated the depletion of muscle ATP levels. The reduction in the cellular energy status during the early postmortem state subsequently leads to the earlier and quicker activation of AMPK, which increases the risk of the development of PSE in pork loin (Shen et al., 2007; Shen et al., 2006).

Since STO-609 injections in murine LL muscle tissue result in decreased glycolysis, they have implications for the regulation of AMPK in this type of tissue. The results of these experiments in postmortem skeletal muscle suggest the partial regulation of glycolysis by AMPK. Previous studies suggest that glycolysis in ischemic cardiac muscle is increased by AMPK at two primary points: one is phosphor-fructose kinase 1 (PFK1), while the other is glycogen phosphorylase. The activation of AMPK can up-regulate glycolysis due to its ability to activate and phosphorylate phosphorylase kinase. In turn, phosphorylated kinase can then phosphorylate and activate glycogen phosphorylase, which is an enzyme controlling glycogenolysis and catalyzing glycolysis substrate production (Russell et al., 1999; Young et al., 1997; Fraser et al., 1999;). In addition, activated AMPK is responsible for the phosphorylation and activation of phosphofructokinase-2 (PFK-2) (Marsin et al., 2000). In turn, activated PFK-2 catalyzes fructose-2,6-phosphate production. PRK-1 is the most important enzyme at controlling the rate of glycolysis, and the enzyme is activated allosterically by fructose-2,6-phosphate.

In humans and mice, AICAR is commonly used as an AMPK activator; its role in AMPK activation in LL muscle was established by McFadden and Corl (2009). This role was confirmed in yak LL muscle during differentiation in this study. As expected, STO-609 injection decreased the phosphorylation of AMPK in LL muscle from postmortem yak, but AICAR has the opposite effect. AMPK could still be activated in postmortem yak skeletal muscle that had not been injected with STO-609, and the greatest amounts of activity were detected in the 12 h postmortem samples. However, the postmortem samples that had been injected with STO-609 did not exhibit any changes in the activity of AMPK activity over time. In addition, the postmortem yak samples that had been treated with STO-609 had lower levels of ACC phosphorylation at Thr 172. This result was consistent with the lower amounts of AMPK activity and also with the findings of previous research (Shen & Du, 2005a; Du et al., 2005). In concert, these findings confirm that STO-609 is a potent inhibitor of AMPK, which leads to lower levels of activity of AMPK in the postmortem skeletal yak muscle.

We implemented real-time RT-PCR to examine AMPK mRNA expression affected by AICAR. The purpose of this experiment was to identify the mechanisms by which AICAR induces the activation of the AMPK protein. Figure 5 shows that the expression of the mRNA of AMPK increased significantly at 12 h after treatment with AICAR. This figure also shows the dose-dependent manner in which 12 h treatments with STO-609 induced effects opposite to those of AICAR. Thus, suppression of AMPK mRNA expression was one of the mechanisms by which STO-609 induced the reduction of the AMPK protein. However, the mechanisms by which AICAR induces a high level regulation of the AMPK gene via AMPK activation remain unclear. One possible mechanism by which AMPK could be transcriptionally regulated could involve the modification of transcription factors by their direct phosphorylation. In addition, it has been reported that AMPK decreases the amount of glucose by modifying its stability. These results raise the possibility that AMPK could directly target glucose and that its phosphorylation of glucose could increase the rate of degradation of this compound. Another way in which AMPK-induced transcriptional reduction could affect the metabolism of the tissue is through the phosphorylation of the cofactors that control the activity of transcription. For example, AMPK causes a reduction in the affinity of p300 for multiple nuclear receptors by phosphorylating it on Ser89. This reaction results in a decrease in the affinity of p300 for multiple nuclear receptors (e.g. thyroid hormone and peroxisome proliferator-activated receptors (PPAR) c and a). However, this study did not enable us to identify the transcription factors that were responsible for regulation increase in the gene expression of AMPK induced by AICAR. Therefore, addition research to identify the transcription factors and cis-elements that are involved in the response to AICAR is merited.

## 5. Conclusions

The changes in beef glycolysis, energy metabolism, AMPK activity and the expression of the AMPK gene (PRKKA1, PRKKA2) in Yushu yak during the postmortem period were measured. The results of this study demonstrate that the expression of the PRKAA1 and PRKAA2 genes and the AMPK activity were subject to AICAR activation and STO-609 inhibition. This suggests that the increased expression of the PRKAA1 and PRKAA2 genes will increase the activity of AMPK. After AMPK is activated, the direct phosphorylation glycolysis pathway increases the glycolysis activity, which promotes the glycolysis process and produces a large amount of lactic acid. This production results in the decrease of energy metabolism in postmortem animal muscles. Therefore, the activity of the yak AMPK increased under hypoxic adaptation, which accelerated glycolysis and metabolism, and more effectively regulated energy production. It laid a foundation for the establishment of the theoretical system of energy metabolism in post-mortem yak meat.

## Acknowledgments

We thank colleagues in the laboratory and our collaborators for their useful suggestions. This work was supported by the China Agriculture Research System (CARS-37), National Natural Science Foundation of China (Grant Nos: 31760482).

## Reference

Birnbaum, M. J. (2005). Activating AMP-activated protein kinase without AMP. Molecular Cells, 19, 289–290.

Blattler, S. M., Rencurel, F., Kaufmann, M. R., & Meyer, U. A. (2007). In the regulation of cytochrome P450 genes, phenobarbital targets LKB1 for necessary activation of AMP-activated protein kinase. Proceedings of the National Academy of Science USA, 104, 1045–1050.

Briskey, E. J., & Wismer-Pedersen, J. (1961). Biochemistry of pork muscle structure. 1. Rate of anerobic glycolysis and temperature change versus apparent structure of muscle tissue. Journal of Food Science, 26, 297–305.

Cannon, J. E., Morgan, J. B., McKeith, F. K., Smith, G. C., Sonka, S., Heavner, J., et al. (1996). Pork chain quality audit and survey: Quantification of pork quality characteristics. Journal of Muscle Foods, 7, 29–44.

Carling, D. (2004). The AMP-activated protein kinase cascade – A unifying system for energy control. Trends in Biochemical Science, 29, 18–24.

Carling, D., Zammit, V. A., & Hardie, D. G. (1987). A common bicyclic protein-kinase cascade inactivates the regulatory enzymes of fatty-acid and cholesterol-biosynthesis. FEBS Letters, 223, 217–222.

Chen Z P, Stephens T J, Murthy S, et al. (2003). Effect of exercise intensity on skeletal muscle AMPK signaling in humans. Diabetes, 52, 2205–2212.

Copenhafer, T. L., Richert, B. T., Schinckel, A. P., Grant, A. L., & Gerrard, D. E. (2006). Augmented postmortem glycolysis does not occur early postmortem in AMPK73-mutated porcine muscle of halothane positive pigs. Meat Science, 73, 590–599.

Corton, J. M., Gillespie, J. G., & Hardie, D. G. (1994). Role of the AMPactivated protein kinase in the cellular stress response. Current Biology, 4, 315–324.

Crute, B. E., Seefeld, K., Gamble, J., Kemp, B. E., & Witters, L. A. (1998). Functional domains of the alpha1 catalytic subunit of the AMP-activated protein kinase. Journal of Biological Chemistry, 273, 35347–35354.

Ding, X. Z, Liang, C. N, Guo, X, et al. (2014) Physiological insight into the high-altitude adaptations in domesticated yaks (*Bos grunniens*) along the Qinghai-Tibetan Plateau altitudinal gradient. Livestock Science, 162, 233–239.

Fan, X., Ding, Y., Brown, S., Zhou, L., Shaw, M., Vella, M.C., Cheng, H., McNay, E.C., Sherwin, R.S., McCrimmon, R.J., (2009). Hypothalamic AMP-activated protein kinase activation with AICAR amplifies counterregulatory responses to hypoglycemia in a rodent model of type 1 diabetes. Am. J. Physiol. Regul. Integr. Comp. Physiol, 296, R1702–R1708.

Hardie, D. G., & Carling, D. (1997). The AMP-activated protein kinase– fuel gauge of the mammalian cell? European Journal of Biochemistry, 246, 259–273.

Hardie, D. G., Salt, I. P., Hawley, S. A., & Davies, S. P. (1999). AMP activated protein kinase: An ultrasensitive system for monitoring cellular energy charge. Biochemical Journal, 338, 717–722.

Hardie, D. G. (2003) Minireview: the AMP-activated protein kinase cascade: the key sensor of cellular energy status. Endocrinology, 144, 5179–5183

Hardie, D. G., Scott, J. W., Pan, D. A., & Hudson, E. R. (2003). Management of cellular energy by the AMP-activated protein kinase system. FEBS Letters, 546, 113–120.

Hardie, D. G. (2004). AMP-activated protein kinase: The guardian of cardiac energy status. Journal of Clinical Investigation, 114, 465–468.

Hardie, D. G., Ross, F. A., Hawley, S. A. (2012). AMPK: a nutrient andenergy sensor that maintains energy homeostasis. Nature Reviews Molecular Cell Biology, 13, 251–262.

Hou, Y., Yao, K., Wang, L., Ding, B., Fu D & Wu, g. (2011).Effects of alpha-ketoglutarate on energy status in the intestinal mucosa of weaned piglets chronically challenged with lipopolysaccharide. British journal of nutrition,106,357–363.

Kemp, B. E., Mitchelhill, K. I., Stapleton, D., Michell, B. J., Chen, Z. P., & Witters, L. A. (1999). Dealing with energy demand: The AMP activated protein kinase. Trends in Biochemical Science, 24, 22–25.

Kim, E. K., Miller, I., Aja, S., Landree, L. E., Pinn, M., McFadden, J., et al. (2004). C75, a fatty acid synthase inhibitor, reduces food intake via hypothalamic AMP-activated protein kinase. Journal of Biological Chemistry, 279, 19970–19976.

Lee, Y. B., & Choi, Y. I. (1999). PSE (pale, soft, exudative) pork: The causes and solutions – Review. Asian Australian Journal of Animal Sciences, 12, 244–252.

Minokoshi, Y., Alquier, T., Furukawa, N., Kim, Y. B., Lee, A., Xue, B., et al. (2004). AMP-kinase regulates food intake by responding to hormonal and nutrient signals in the hypothalamus. Nature, 428, 569–574.

Monin, G., & Sellier, P. (1985). Pork of low technological quality with a normal rate of muscle pH fall in the immediate post-mortem period: The case of the Hampshire breed. Meat Science, 13, 49–63.

Musi, N, Hayashi, T, Fujii, N, et al. (2001). AMP-activated protein kinase activity and glucose uptake in rat skeletal muscle. Am J Physiol Endocrinol Metab, 280, E677–E684.

Park, S. H., Gammon, S. R, Knippers J D., et al. (2002). Phosphorylation activity relationships of AMPK and acetyl-CoA carboxylase in muscle. Journal of Appl Physiol, 92, 2475–2482.

Pold, R., Jensen, L.S., Jessen, N., Buhl, E.S., Schmitz, O., Flyvbjerg, A., Fujii, N., Goodyear, L.J., Gotfredsen, C.F., Brand, C.L., Lund, S. (2005). Long-term AICAR administration and exercise prevents diabetes in ZDF rats. Diabetes, 54, 928–934.

Proszkowiec-Weglarz, M., Richards, M. P., Ramachandran, R., et al. (2006). Characterization of the AMP-activated protein kinase pathway in chickens. Comparative Biochemistry & Physiology Part B Biochemistry & Molecular Biology, 143, 92–106.

Sambandam, N., & Lopaschuk, G. D. (2003). AMP-activated protein kinase (AMPK) control of fatty acid and glucose metabolism in the ischemic heart. Progress in Lipid Research, 42, 238–256.

Schwagele, F., Lopez, P., Haschke, C., & Honikel, K. O. (1994). Rapid pH drop in PSE-muscles – Enzymological investigations into the causes. Fleischwirtschaft, 74, 95–101.

Shen, Q. W., & Du, M. (2005a). Effects of dietary alpha-lipoic acid on glycolysis of postmortem muscle. Meat Science, 71, 306–311.

Shen, Q. W., & Du, M. (2005b). Role of AMP-activated protein kinase in the glycolysis of postmortem muscle. Journal of Science of Food Agriculture, 85, 2401–2406.

Shen, Q. W., Jones, C. S., Kalchayanand, N., Zhu, M. J., & Du, M. (2005). Effect of dietary alpha-lipoic acid on growth, body composition, muscle pH, and AMP-activated protein kinase phosphorylation in mice. Journal of Animal Science, 83, 2611–2617.

Shen, Q. W., Means, W. J., Thompson, S. A., Underwood, K. R., Zhu, M. J., McCormick, R. J., et al. (2006). Pre-slaughter transport, AMP-activated protein kinase, glycolysis, and quality of pork loin. Meat Science, 74, 388–395.

Shen, Q. W. W., Means, W. J., Underwood, K. R., Thompson, S. A., Zhu, M. J., McCormick, R. J., et al. (2006). Early post-mortem AMPactivated protein kinase (AMPK) activation leads to phosphofructokinase-2 and-1 (PFK-2 and PFK-1) phosphorylation and the development of pale, soft, and exudative (PSE) conditions in porcine longissimus muscle. Journal of Agricultural and Food Chemistry, 54, 5583–5589.

Shen, Q. W., Underwood, K. R., Means, W. J., McCormick, R. J., & Du, M. (2007). The halothane gene, energy metabolism, adenosine monophosphate-activated protein kinase, and glycolysis in postmortem pig longissimus dorsi muscle. Journal of Animal Science, 85, 1054–1061.

Sieczkowska, H., Koc’win-Podsiadła, M., Zybert, A., Krzecio, E., Antosik, K., Kamin’ski, S., Wójcik, E. (2010). The association between polymorphism of PKM2 gene and glycolytic potential and pork meat quality. Meat Science, 84, 180–185.

Solomon, M. B., Van Laack, R. L. J. M., & Eastridge, J. S. (1998). Biophysical basis of pale, soft, exudative (PSE) pork and poultry muscle: A review. Journal of Muscle Foods, 9, 1–11.

Stein, S. C., Woods, A., Jones, N. A., Davison, M. D., & Carling, D. (2000). The regulation of AMP-activated protein kinase by phosphorylation. Biochemical Journal, 345, 437–443.

Thomson, D. M., Brown, J. D., Fillmore, N., Ellsworth, S. K., Jacobs, D. L., Winder, W. W., Fick, C. A., Gordon, S. E., (2009). AMP-activated protein kinase response to contractions and treatment with the AMPK activator AICAR in young adult and old skeletal muscle. Journal of Physiol, 587, 2077–2086.

Vincent, M. F., Erion, M. D., Gruber, H. E., and Van den Berghe, G. (1996) Hypoglycaemic effect of AICA riboside in mice. Diabetologia, 39, 1148–1155.

Winder, W. W. (2001). Energy-sensing and signaling by AMP-activated protein kinase in skeletal muscle. Journal of Applied Physiology, 91, 1017–1028.

Woelfel, R. L., Owens, C. M., Hirschler, E. M., Martinez-Dawson, R., & Sams, A. R. (2002). The characterization and incidence of pale, soft, and exudative broiler meat in a commercial processing plant. Poultry Science, 81, 579–584.

Wojtaszewski, J. F., Nielsen, P., Hansen, B. F., et al. (2000). Isoform specific and exercise intensity-dependent activation of 5′-AMP-activated protein kinase in human skeletal muscle. Journal of Physiol, 528, 221–226.

Woods, A., Johnstone, S. R., Dickerson, K., Leiper, F. C., Fryer, L. G. D., Neumann, D., et al. (2003). LKB1 is the upstream kinase in the AMP-activated protein kinase cascade. Current Biology, 13, 2004–2008.

Zuo, H. X., Han, L., Yu, Q., et al. (2017) Proteomics and bioinformatics analyses of differentially expressed proteins in yak and beef cattle muscle. Transactions of the Chinese Society for Agricultural Machinery, 48, 313–320.

